# Partitioning the genomic journey to becoming *Homo sapiens*

**DOI:** 10.1101/2024.12.09.627480

**Authors:** Luca Pagani, Riccardo Bertazzon, Vasili Pankratov, Leonardo Vallini, Davide Marnetto, Adeline Morez, Hélios Delbrassine, Francesca Carollo, Maurizio D’Esposito, Ilaria Granata, En En Teo, Aswini Leela Loganathan, Pille Hallast, Charles Lee, Qasim Ayub, Massimo Mezzavilla

## Abstract

What makes us human? *Homo sapiens* diverged from its ancestors in fundamental ways, as reflected in recent genomic acquisitions, such as the PAR2-Y chromosome translocation. Here we show that despite morphological and cultural differences between modern and archaic humans, these human groups share these recent acquisitions. Our modern lineage shows recent functional variants in only 56 genes, of which 24 are linked to brain functions and skull morphology. Nevertheless, these acquisitions failed to introgress into Neanderthals when archaic and modern populations admixed after 350 kya, suggesting their exclusive link to the modern human niche or that Neanderthal’s small population size hindered their spread. Taken together, our results point to a scenario where Modern and Archaic should be regarded as populations of an otherwise common human species, which independently accumulated mutations and cultural innovations.

## Introduction

The acquisition of derived traits has punctuated the long evolutionary path followed by the ancestors of all humans, as witnessed by the fossil record. This constant accumulation of physical, cognitive and cultural innovations is what we collectively refer to when we think of ourselves as “humans”. The emerging consensus, corroborated by fossil ^1,2,3–5^ and molecular ^6–8^ evidence, sees the emergence of modern humans from an evolutionary branch that split from that of Neanderthal and Denisovans (collectively referred to as archaic humans) around 650 kya^7^ (from now on labelled as “Split” ). Recent molecular evidence added complexity to this simple split scenario, positing the existence of a deep structure within the human lineage, namely population A and B, with a separation dated to around 1.5 Mya ^9^. Population A would be responsible for the entire gene pool of Neanderthals and Denisovans and for 80% of the gene pool of modern humans, while population B would have contributed up to 20% to modern humans, rejoining the main population A about 300 kya. Interestingly, the A/B population structure merger took place before the deepest reported population splits between modern human populations^8^, hence featuring among the last major genomic events of our history as a species. As the evolutionary period extending from the separation of modern and archaic human lineages to the earliest splits between modern human populations remains elusive, one may wonder whether it promoted the accumulation of modern human derived elements, or whether such elements were already available before the separation from archaics, rendering, de facto, modern and archaic humans physiologically “human”. ^10^.

In recent years, growing evidence has accumulated in favour of an admixture event between an early modern human lineage and Neanderthals about 350 kya, as witnessed by uniparental (mtDNA and chrY) ^11,12,13,14,15^ and genomic evidence ^12,16–18^. Importantly, this latter introgression occurred as early as 300 ky prior^19^ and in the opposite direction with respect to the more widely known human-Neanderthal introgression^20^ that followed the Out of Africa expansion of the modern human population. It thus provides an excellent time checkpoint to assess whether the genetic substitutions, accumulated along the modern human lineage after the separation from archaics (Split, **Figure 1A** in dark violet), displayed a high enough adaptive potential to be introgressed and then fixed in the Neanderthal gene pool (**Figure 1A**, in blue). If so, one may postulate that such mutations could yield, at least in part, the answer to what defines us as modern humans. Otherwise, one should conclude that all key genetic substitutions were already present before the Split and were hence shared by modern and archaic humans alike, or that they emerged as a set of modern human-specific substitutions after the Introgression (either from *de novo* mutation or inherited from population B during the population A/B merger) (**Figure 1A**, in red).

**Figure 1.**
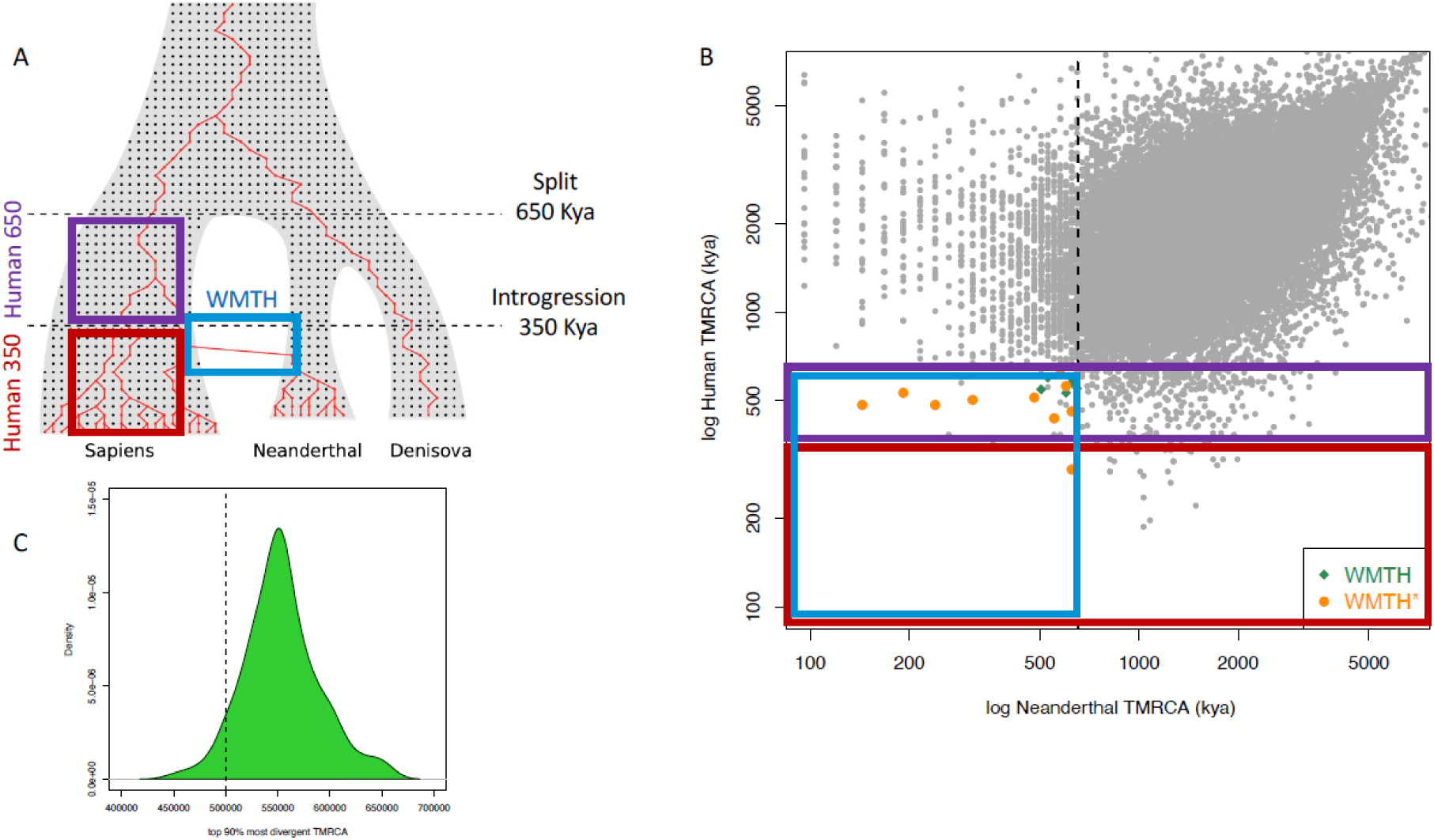
A. Events Partitioning. Defining the Split (the separation of modern and archaic lineages, around 650 kya) and the Introgression (the contribution of modern human lineages to the gene pool of Neanderthals, after 350 kya) along the population tree of modern and archaic humans. The gene tree depicted in red exemplifies a Human650 region that entered and was also fixed within the Neanderthal gene pool as a consequence of the Introgression.. B )Scatterplot of the Neanderthal-Human reference ( log x-axis) and the Human-Human ( log y-axis) TMRCAs on windows displaying a Denisova-Human reference TMRCA >=650 kya. Grey dots represent the bulk of the distribution, which, as expected, is centered around the diagonal for x and y values >=650 kya. Dashed lines report the date of the Split (the separation of modern and archaic humans) along both axes.The top left quadrant is, as expected, enriched by Neanderthal regions introgressed inside the reference sequence, but we assume this signal not to impact our results, since the resulting human-human TMRCA (y-axis) is above the thresholds used for our downstream analyses. Orange dots and green diamonds are the 50 kb regions reported in Table 1 with more (orange) and less (green) stringent criteria (What made them humans, WMTH regions). C) Density plot of the top 90% most divergent TMRCA estimates obtained for every 50-kb window along our 5 Mbp-long simulated genomes. The dotted line indicates the real TMRCA simulated (500,000 years ago).

**Table 1.** “What made them humans” (WMTH) regions. 17 Human650 regions that contributed to the gene pool of Neanderthals as a consequence of the Introgression event. The last seven regions, highlighted with an asterisk, display at least one derived site shared between modern humans and Denisova (D-Hder) but not Neanderthal and hence do not meet our full list of inclusion criteria. The abundance of regions landing on chromosome X is explained by the smaller effective population size of its gene pool, which in turn affects the distribution of expected neutral coalescences.

| Chr | Start(kb) | End(kb) | NeaTMRCa | DenTMRCa | HumTMRCa | Genes | N-Hder(R,C) | D-Hder(R,C) | Hder(R,C) |
| --- | --- | --- | --- | --- | --- | --- | --- | --- | --- |
| 5 | 45600 | 45650 | 551834 | 743851 | 436800 | HCN1 | 1(0,0) | 0(0,0) | 4(0,0) |
| 6 | 27900 | 27950 | 239976 | 3791166 | 484800 | OR2W6P;RPLP2P1;OR2W4P;OR2B6 | 47(0,0) | 0(0,0) | 0(0,0) |
| 7 | 30650 | 30700 | 143954 | 935888 | 484800 | GARS;CRHR2 | 1(0,0) | 0(0,0) | 1(0,0) |
| 10 | 103600 | 103650 | 623763 | 815804 | 460800 | KCNIP2;C10orf76 | 5(1,0) | 0(0,0) | 0(0,0) |
| 10 | 103650 | 103700 | 599820 | 791762 | 561600 | C10orf76 | 8(0,0) | 0(0,0) | 4(0,0) |
| 12 | 123550 | 123600 | 575896 | 1127774 | 643200 | RP11-197N18.7;PITPNM2 | 4(0,0) | 0(0,0) | 1(0,0) |
| 16 | 22000 | 22050 | 191946 | 1031670 | 532800 | CTD-2649C14.1;PDZD9;C16orf52 | 12(0,0) | 0(0,0) | 0(0,0) |
| 16 | 22050 | 22100 | 311919 | 1223706 | 504000 | RP11-101E7.2;C16orf52 | 8(0,0) | 0(0,0) | 0(0,0) |
| X | 25350 | 25400 | 479952 | 1007798 | 513600 | - | 4(0,0) | 0(0,0) | 7(0,0) |
| X | 62950 | 63000 | 623925 | 671906 | 292800 | ARHGEF9 | 1(0,0) | 0(0,0) | 1(0,0) |
| 3* | 51650 | 51700 | 647857 | 791842 | 552000 | RAD54L2;TEX264 | 2(0,0) | **2(0,0)** | 2(0,1) |
| 3* | 99850 | 99900 | 527873 | 1487613 | 600000 | RP11-15N24.4;CMSS1 | 13(0,1) | **1(0,0)** | 0(0,0) |
| 11* | 118300 | 118350 | 599796 | 863862 | 532800 | RN7SL86P;ATP5L;RP11-770J1.4;KMT2A | 1(0,0) | **1(0,0)** | 0(0,0) |
| X* | 81400 | 81450 | 623900 | 2135402 | 576000 | - | 26(0,0) | **2(0,0)** | 1(0,0) |
| X* | 81500 | 81550 | 503859 | 1511426 | 547200 | - | 10(0,0) | **1(0,0)** | 7(0,0) |
| X* | 96650 | 96700 | 647832 | 1319525 | 374400 | DIAPH2 | 10(0,0) | **2(0,0)** | 1(0,0) |
| X* | 119650 | 119700 | 623888 | 1439741 | 600000 | CUL4B | 14(0,0) | **1(0,0)** | 0(0,0) |

**Abbreviations:** chr: chromosome; NeaTMRCa: time to the most recent common ancestor (TMRCa) between the Altai Neanderthal sequence and the hg19 human reference; DenTMRCa: TMRCa between the Denisova sequence and the hg19 human reference; HumTMRCa: top 90% deepest TMRCa between a panel of 50 African human genomes and the hg19 human reference; N-Hder: sites that are homozygous derived in Altai Neanderthal and have a derived allele frequency of at least 98% in the 1000 Genomes Samples individuals and are homozygous ancestral in Denisova; R,C: of these, how many sites are annotated with Regulatory or Coding consequences (See Table S2 for a full list of these sites); D-Hder: sites that are homozygous derived in Denisova and have a derived allele frequency of at least 98% in the 1000 Genomes Samples individuals and are homozygous ancestral in Altai Neanderthal. All genomic coordinates are in build37 (hg19).

With this work, we set out to exploit the Split and Introgression events to partition human-specific genomic loci ^21^ and ultimately shed light on the long-lasting question of what makes us human.

## Results

### Partitioning the genomic landscape of modern and archaic humans

We used the high coverage genomes of Altai Neanderthal and Denisova (Denisova3) as representatives of archaic humans, and of 50 Sub-Saharan individuals from the 1000 Genomes Project as representatives of modern humans. For each individual, we calculated the Time to Most Recent Common Ancestor (TMRCA) with the human reference genome (treated as a random human individual) in 50kbp long windows along the genome (**Figure 1 B, Table S1**).

TMRCA estimates were based on pairwise sequence differences and assumed an average genomic mutation rate of 1.25 * 10-8 mut/bp/gen.. We show with coalescent simulations that this simple approach recapitulates the true TMRCA when the top 90% most divergent TMRCA is taken as a summary statistic when 50 modern humans are considered (**Figure 1C**). We excluded from downstream analyses genomic windows with a human-Denisova TMRCA younger than 650 kya, considered enriched in alterations of the local molecular clock (**Figure 1A**).

We then focused on genomic windows that showed a TMRCA among modern humans comprised between 650 kya (Split) and 350 kya (Introgression), or more recently than 350 kya (Human650 and Human350 regions, respectively, see Methods and **Figure 1).** This second window was chosen for its potential of having experienced fixation events in modern humans after the Introgression. Of all the genomic windows displaying at least one missense variant with derived frequency >=98% in modern humans and fixed derived in both Altai Neanderthal and Denisova (Figure S1, golden diamond) ^22^, only a 16 are indeed tagging recent events (Human 650 and Human 350) along the modern human lineage and can be deemed as highly informative in defining what makes us human, additionally showing the high specificity of our approach.

In addition, we were also interested in how many regions showing a recent human TMRCA (Human650 and Human350) were transferred to the Neanderthal gene pool as a consequence of Introgression, hence displaying versatile and high adaptive potential also in this population of archaic humans. As expected, given the “Introgression” event, we show that Altai Neanderthal displays more regions with a recent coalescence event (<= 650 kya) with the human reference genome than Denisova **(Figure S2,** blue line). This pattern may also be affected by the widespread introgression of Neanderthal alleles into Eurasian genomes ^20^, including into a non-negligible portion of the hg19 reference genome used for these analyses (as also visible in the top left quadrant of **Figure 1B and S1**). However, an even more striking difference between the Neanderthal and Denisova signals, which can only be explained as a consequence of the “Introgression”, is observed when looking only within Human650 regions (**Figure S2,** green line), which show an excess of reference-Neanderthal recent coalescences compared to reference-Denisova, maxing when the Altai Neanderthal-reference human genome coalescences are around the time of Introgression. Furthermore, within the Human650 regions, the number of regions with a Neanderthal-reference human genome coalescence <= 650 kya (Neanderthal Recent) (n=22) is significantly higher (Fisher exact test, p-value 0.01) than the number of regions (n=7) expected if Human650 and Neanderthal Recent were two independent sets of regions. This result points to the Introgression event having indeed caused a significant number of Human650 regions to have entered the Neanderthal gene pool.

We then tried to single out the Human650 genomic regions that entered the Neanderthal gene pool extracting those with 1) a human reference-Neanderthal coalescence <=650kya, 2) at least one site with derived allele frequency >=98% in modern humans, homozygous derived in Altai Neanderthal and homozygous ancestral in Denisova and 3) no sites with derived allele frequency >=98% in modern humans, homozygous derived in Denisova and homozygous ancestral in Altai Neanderthal. We dubbed these regions “what made them humans” (WMTH), being Human650 regions that contributed to their (Neanderthal’s) gene pool. We found ten WMTH regions (**Table 1**, first ten rows, **Figure 1B and S1**, orange dots) that matched all three criteria and an additional seven that matched just the first two criteria (**Table 1**, bottom seven rows marked by an asterisk, **Figure 1B and S1**, green diamonds). Many such regions land on chromosome X, since its smaller population size compared to the autosomes is responsible for a higher coalescence rate. Further validation of the inferred TMRCA of the WMTH regions was performed with Relate (**Supplementary Information, Figure S3**). We also report one region on chromosome X (X:62950000-63000000) that met the WMTH criteria albeit belonging to the Human350 category. We decided to include it in the analysis to avoid spurious threshold effects to be introduced by our decision to partition human regions in Human650 and Human350 mutually exclusive categories.

### Functional characterization

Of all the derived alleles shared between Neanderthal and modern humans in WMTH regions, only two (**Table 1**, in bold) were predicted to have regulatory or coding consequences, with one of the two falling in a region not meeting our 3rd inclusion criterion. We asked whether this apparent scarcity of functional mutations is expected when looking at the broader sets of Human650 (**Figure 2**). Overall, WMTH and Human650 regions encompass a similar proportion of functional sites (4.35% and 4.28%, respectively) and the two sets are similar also when looking at the functional sites shared by modern human and Neanderthals, but not Denisovans (FuncHN, Figure 2) (2 regions with at least 1 FuncHN out of 11 WMTH regions (18.2%) versus 60 out of 242 Human650 regions, 24.7%.

**Fig 2.**
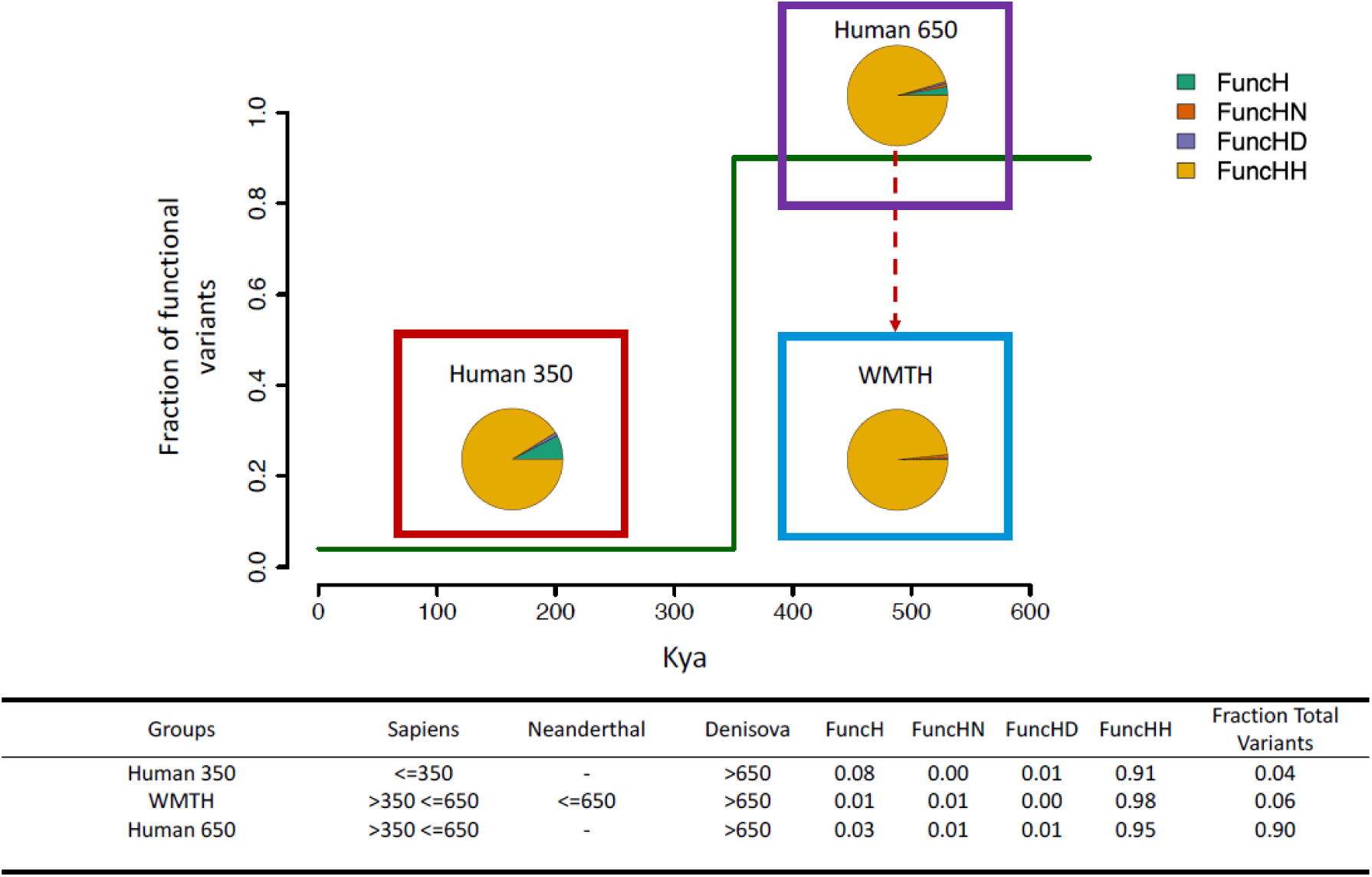
Prevalence of lineage-specific functional variants according to the modern human TMRCA (shown on the x-axis) of the 50 kb window. The green line represents the relative fraction of fixed-derived human functional variants exhibited by the genomic windows showing modern human TMRCA <= 1000 kya. Within each time period, a pie chart represents the breakdown of functional derived sites into FuncH (shared by >98% of modern humans but no archaics; FuncHN, shared by >=98% of modern humans, homozygous derived in Altai Neanderthal and homozygous ancestral in Denisova; FuncHD, shared by >=98% of modern humans, homozygous derived in Denisova and homozygous ancestral in Altai Neanderthal; FuncHH, shared by >=98% of modern humans and homozygous derived in both Denisova and Altai Neanderthal. Most functional variants falling in a Human650 region with a TMRCA in modern humans <= Introgression (Human 350) are shared with all three lineages and only a small portion is unique to modern humans (FuncH). Among the variants with a TMRCA in modern humans between Split and Introgression (Human650), we observe that only a negligible fraction is FuncHN. This fraction is significantly reduced in the WMTH subset (i.e., variants that entered the Neanderthal gene pool during the Introgression).The WMTH subset, on the other hand,shows an enrichment in FuncHH variants.

Fisher exact test p-value=1). We then characterized the genomic sites that entered the Neanderthal pool during Introgression by computing the ratio of FuncHN against the sum of functional sites where just modern humans (FuncH), modern humans and Neanderthals (FuncHN), or modern humans and Denisovans (FuncHD) shared the derived allele (e.g., all the sites that were derived in modern humans and that could enter the Neanderthal pool during Introgression). The Neanderthal gene pool received significantly fewer modern human-derived functional sites than it could have, given the ones available at the time of Introgression (2 FuncHN out of 177 FuncHN, FuncHD, FuncH or FuncHH in WMTH regions, or 1.13%, compared to 112 FuncH, FuncHD or FuncHN out of 2465 FuncH, FuncHD, FuncHN or FuncHH in Human650, or 4.5%. Fisher exact test p-value=0.03). It follows that, in contrast, WMTH regions showed an excess of derived sites shared by all archaic and modern humans (FuncHH, in gold in Figure 3), hence having occurred along the lineage shared by all archaic and modern humans,here dubbed the *Homo heidelbergensis* (*HH*) lineage. In other words, WMTH regions, a subset of Human650 regions, are not enriched in modern human-derived functional sites. Conversely, and in line with what was observed for human uniparental markers, which were shown to have permeated the gene pool of Neanderthals because of the accumulation of deleterious variants on the latter ^15^, our results based on autosomal and chromosome X data confirm that Introgression consequences on the Neanderthal gene pool can be best described as a reintroduction of functional variants that were lost in Neanderthal due to the strong genetic drift caused by their reduced effective population size, rather than as an introduction of novel, modern human-specific derived sites.

**Fig 3.**
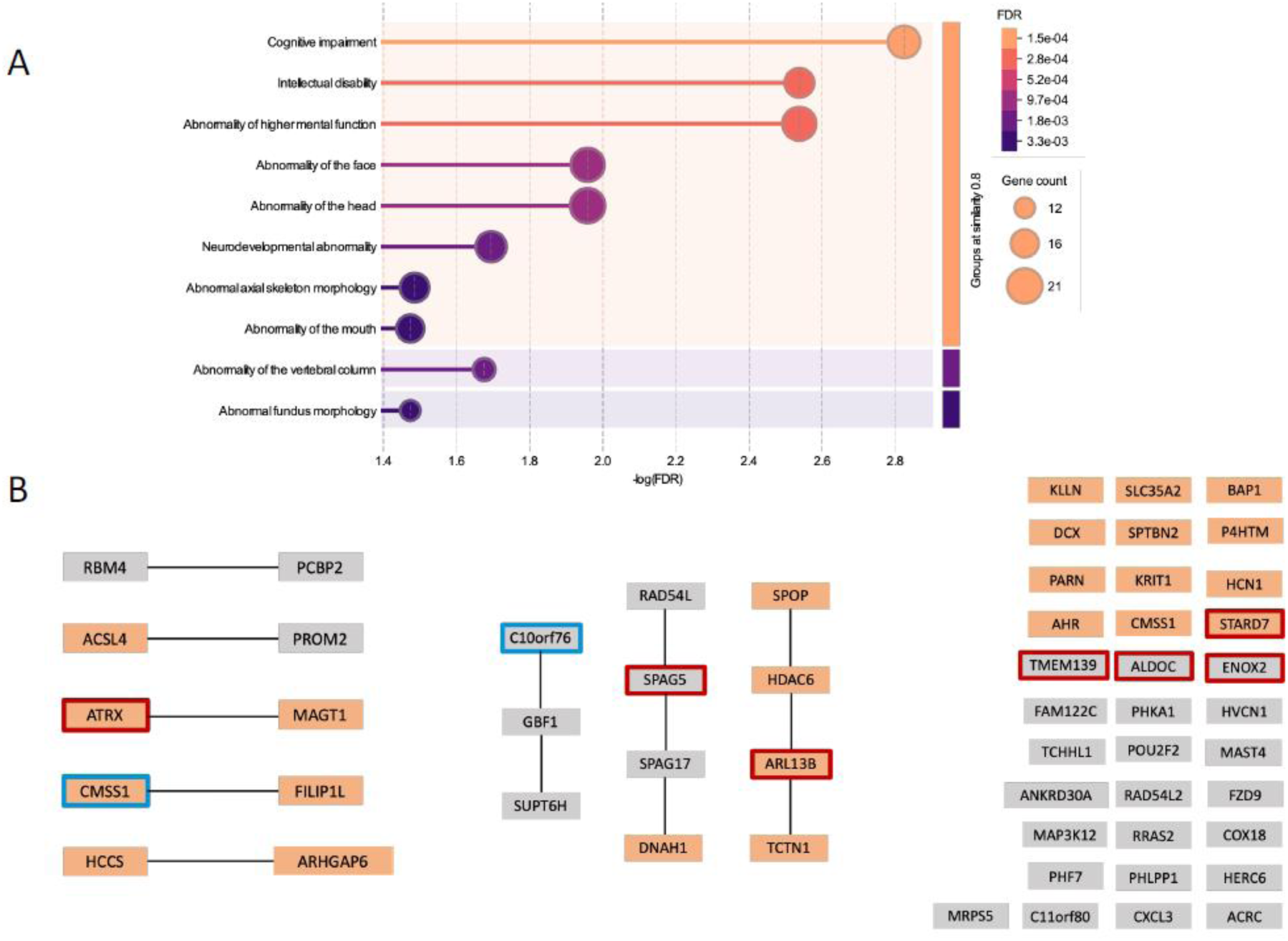
Functional enrichment analyses. A) Functional enrichments of the 56 genes listed in Table S2. Using the STRING database we selected only the phenotypes with a False Discovery Rate (FDR)<=0.05 present in the Monarch phenotypes initiative. The phenotype with the lowest FDR and highest enrichment is cognitive impairment. B) Interaction network obtained from the STRING database using the 56 genes listed in Table S2. Proteins circled in red represent genes with a recent coalescence in humans (Human 350) and proteins circled in blue represent genes defined as WMTH in our manuscript. The 24 orange boxes represent proteins that contribute to at least one of the phenotype enrichments described in panel A.

**Fig 4.**
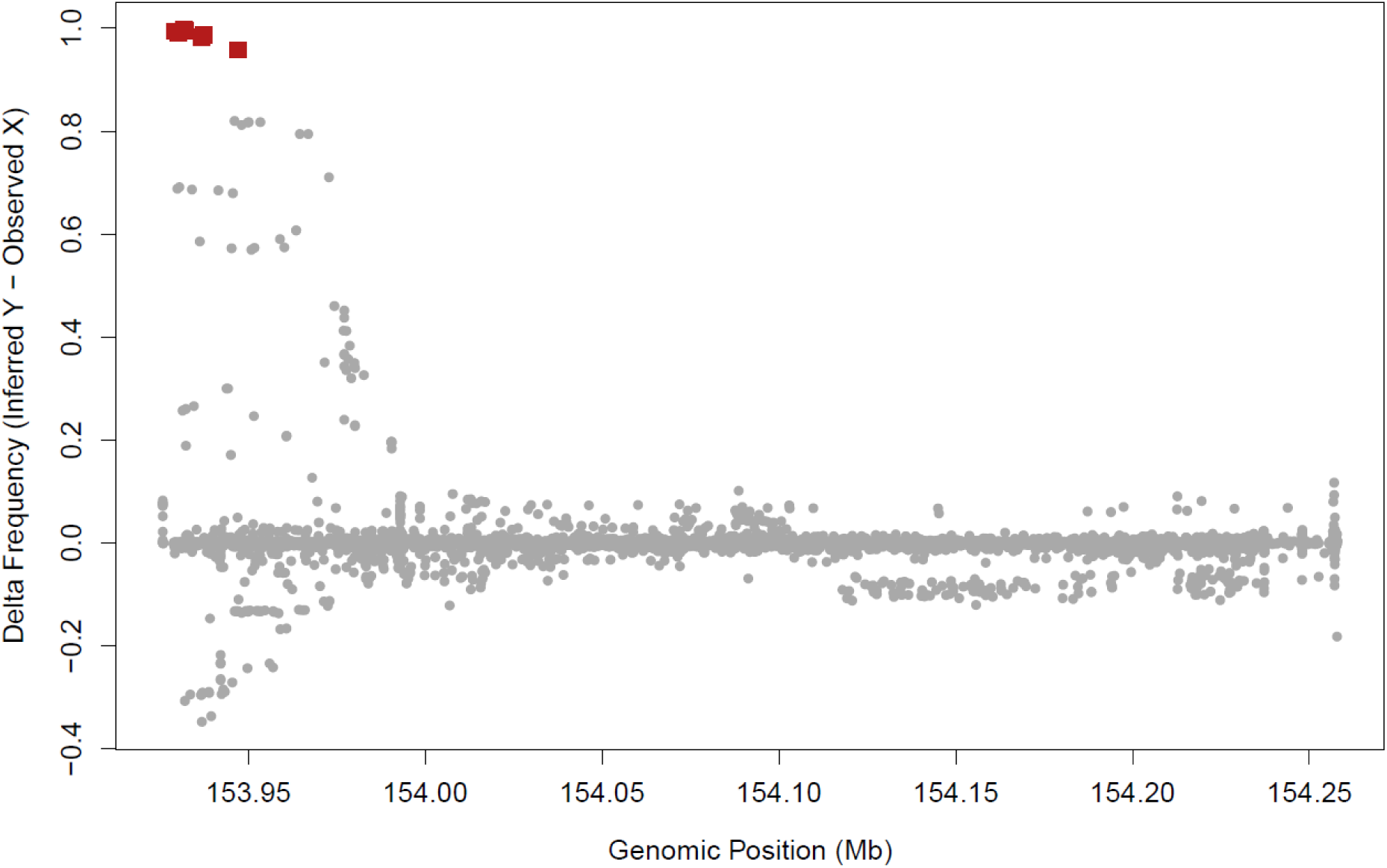
PAR2 Y-specific mutations. Red squares mark the location of the eleven male-specific variants along the ∼330 kb long PAR2 sequence when reads from both chromosomes of modern humans are collapsed together. (Red SNP positions: 153929157; 153929840; 153929878; 153930185; 153931728; 153931814; 153932007; 153932008; 153936764; 153937415; 153947049, mapped onto X-PAR2 with T2T-CHM13v2.0 coordinates, with the PAR2 starting point set at 153925834).

An interpretation of this emerging picture may imply that the vast majority of modern human-derived variants that experienced a human sweep between Split and Introgression encompassed an adaptive potential that contributed perhaps to the biological characterization of our lineage but that, on the other hand, was either not sufficiently overwhelming to swarm into the Neanderthal gene pool or linked to specific environmental or cultural features of the modern human niche. To better understand the biological significance of the variants enclosed within Human650 and Human350 regions, we generated protein-protein interaction networks based on genes falling within a Human650 or Human350 region and displaying a coding or regulatory human-derived variant (here defined as being fixed derived in at least 98% of the 1000 Genomes Project individuals) (**Fig 3**, **Table S2**, **Fig S1**). The resulting list of 56 proteins (**Fig 3)** are, as a whole, significantly enriched in a compelling list of phenotypes (**Fig 3 A**)^23^: the first three (Cognitive impairment, Intellectual disability, Abnormality of higher mental function) are related to higher-level brain function, while the next three (Abnormality of the face, Abnormality of the head, Neurodevelopmental abnormality) are related to developmental anomalies. The 24 proteins associated with one or more of them are highlighted in orange in the network below. Despite these phenotypes appearing to be pivotal aspects to becoming modern humans, only two of these genes (*CMSS1* and *C10orf76*, circled in blue) were passed on to Neanderthals during the Introgression (WMTH regions) and only *CMSS1* is both involved in the above-listed phenotypes and associated with a moderate CADD score (Combined Annotated Dependent Depletion, a measure of the deleteriousness of a mutation)^24^ (CADD=3.27, **Table S2**). On the other hand, seven genes involved in the network (*ATRX, SPAG5, ARL13B, TMEM139, ALDOC, STARD7* and *ENOX2*) experienced a human sweep only after Introgression (**Fig S1**, cyan dots), meaning that their functional variants were potentially not available to Neanderthals during the Introgression and for sure did not enter their gene pool. Notably, SPAG5, ATRX and ARL13B interact with and expand the network of proteins for which derived functional variants were already available at the time of the Introgression (**Fig 3 B**). A tempting explanation may involve the role of these latter three genes as the “detonator” to unlock the biological adaptive potential of other genes (namely *MAGT1, RAD54L, SPAG17, DNAH1, SPOP, HDAC*6 and *TCTN1*), that were not sufficient on their own to trigger a sweep in the Neanderthal gene pool after the Introgression.

Importantly, *SPAG5* features in several published lists as the gene with the highest number of missense substitutions between modern humans and Neanderthals, and its function has been previously associated with the mitotic spindle ^2525,26^. Additional evidence in support of the biological relevance of the modern human-derived allele for the reported SNPs (**Table S2**) is provided by the observation that the vast majority of the functional SNPs falling in the Human650 and Human350 regions display 0% frequency of the ancestral and archaic (Neanderthal or Denisova) allele, even in populations such as Papuan Highlanders ^27,28^, who experienced introgressions from both populations of archaic humans (**Table S2**).

### Newly observed chromosomal rearrangement

Another probable functional consequence of the Introgression is the transfer to the Neanderthal pool, along with the modern human chromosome Y, of the human pseudo-autosomal region 2 (PAR2). Modern human PAR2, like the chromosomal fusion that generated the human chromosome 2, is a genomic rearrangement (X to Y translocation) known to have emerged after the separation of our lineage from the chimpanzee lineage^29^. Together with PAR1, PAR2 enables homologous pairing of the X and Y sex chromosomes during meiosis in human males, but the timing of its translocation event is currently unknown. Firstly, we observed that the male Denisova genome displayed a sequencing depth on regions matching the X chromosome PAR2 higher than the rest of the chromosome (**Supplementary Figure 4**), although not as high (2-fold) as expected. This result may suggest the presence of PAR2 in the Y chromosome of the ancestors of modern and archaic humans unless recurrent acquisition is contemplated^30^, but further tests are needed to overcome limitations presented by the extremely low sequencing coverage available for this Denisovan male individual. Similar results obtained for a male Neanderthal genome (**Supplementary Figure 4**) are provided as a control, given that Neanderthal Y chromosome replacement is among the known consequences of the Introgression, and the presence of PAR2 in these archaic populations is therefore amply expected. As PAR1 is present among all modern and archaic human Y chromosomes, given its shared origin with the chimpanzee lineage, we use it as further positive control for our analysis. Once again, we note a comparably higher sequencing depth displayed by all modern and archaic humans at this site (**Supplementary Figure 4)** when compared to the rest of chromosome X, although coverage uncertainties due to the low quality of available ancient DNA are evident.

We then set out to provide a molecular dating for the PAR2 translocation and inspected its mutational landscape to characterize the substitutions that occurred since the common ancestor of the PAR2-X sequence that recombined onto chromosome Y. We found 11 SNPs present as heterozygous in male PAR2 chromosome X regions and virtually absent in females. These mutations were further attested in Y chromosomes sequenced using long-read platforms^31^ their appearance after the translocation event, on the male-specific PAR2 lineage. The identified 11 mutations span a 21,215 bp region at the 5’ end of the Y-PAR2 and allowed us to estimate 519 Ky as the time elapsed between i) the separation of the ancestor of all Y-PAR2 sequences from the rest of the X-PAR2 gene pool and ii) the most recent common ancestor of all modern human Y chromosomes, using a Y chromosome mutation rate of 3 x 10^-8^ mut/bp/gen^32^. To this estimate, we must add 338 Ky as the time to the MRCA of all modern human Y chromosomes^33^ to provide the full time depth of the translocation event. This region of relatively clear phylogenetic signal is then followed towards the 3’ telomeric end of the region by additional mutations with a less polarized frequency difference between males and females, as expected under a scenario of increasing recombination events between the X and Y-PAR2 regions as a function of the genetic distance from the translocation point.

Thus, the X->Y PAR2 translocation event must have occurred not earlier than 856 kya (95% CI: 633-1253 kya), predating the separation of modern and archaic human lineages. Although the identified polymorphisms likely introduced reference bias, leading to incorrect read mapping and reduced coverage of the PAR regions in the archaic genomes, we could further confirm that the PAR2-Y sites in Spy Neanderthal and Denisova8 shared the expected number of mutations given their known relationship with the modern human Y chromosome and available coverage <u>(Petr et al. 2020)</u>.

## Discussion

By integrating the genetic evidence in an evolutionary informed manner, we can organize the derived traits that define us as modern humans in a coherent framework. First, we show that regions that underwent a modern human-specific sweep after the separation from archaic humans (Human650 regions) include at least 24 genes involved in higher-level brain function and morphological development, encompassing coding and regulatory variants. Among the Human650 regions, none were involved in reproduction or fertility pathways, suggesting that gametic isolation was not the primary driver of modern and archaic human divergence.

, we can then investigate whether the derived variants within the Human650 regions enhanced fitness specifically the social and ecological niches of modern humans, or whether they might also have benefited Neanderthals. This question can be addressed by considering the Introgression event (the human-Neanderthal admixture occurred after 350 kya), when these variants had the opportunity to enter the Neanderthal gene pool. We found 17 Human650 regions that entered the Neanderthal gene pool (WMTH regions). However, they are not enriched in functional variants and only include 1 of the 24 genes linked with the brain phenotypes mentioned above (circled in blue in Figure 3A). Finally, when focusing on highly deleterious SNPs (CADD score>=20) within Human650 regions, we found rs150596428, a missense in SPAG17, rs4413373, a stop gain in HERC6, and rs1782348480, a missense in AHR. None of them fell within the WMTH regions, and only one of them, AHR, was involved in the brain-related phenotype enrichment highlighted in Figure 3. Additionally, the Human650 genomic regions are instead enriched in functional variants that had already emerged before the Split event (modern, Denisovan and neanderthal divergence). These functional alleles were likely lost by genetic drift in the Neanderthal lineage enhanced by their low effective population size. Of note, this affected the OR2B6, GARS, CRHR2, C10orf76, PDZD9 protein-coding genes, among others (see **Supplementary Table S3** for an inclusive list). Our results suggest that the lost functional variants were not an absolute advantage for fitness, since they were not recovered by Neanderthal through introgression with modern humans.

We additionally identify modern human-specific sweeps having occurred after the Introgression, which encompass the *SPAG5, ATRX* and *ARL13*B genes. Previous studies suggested secondary gene flows from modern humans into Neanderthals between 100 kya ^16^ and 200 kya^12^. Regardless of the exact timing of gene flow, we here show that the modern human haplotypes of *SPAG5, ATRX* and *ARL13*B did not enter the Neanderthal lineage at a later stage, despite their coalescences in modern humans all precede 200 kya and generally pre-date the deepest separations between present-day modern human populations^8,12^.

Overall, whether or not the human-derived variants in *SPAG5, ATRX* and *ARL13B* ever had the opportunity to express their adaptive potential in Neanderthals, the scenario emerging from this work suggests that modern human-derived functional variants that accumulated over the past 650 kya, particularly those linked to higher brain functions, were most likely tightly linked to the ecological and cultural niche occupied by modern humans in Africa.

This notwithstanding, further functional studies are needed to elucidate the putative role of *ATRX, SPAG5* and *ARL13B* as recently acquired keystones of the biological pathways selected by the ecological niche of our human population over the past 650 ky. Indeed, they may have played a role in the fine-tuning of higher-level brain functions, perhaps the most iconic asset of our modern human phenotype.

We also caution that our results build on the explicit assumption that modern human defining features must have experienced a sweep after the Split. This strict experimental framework may overlook additional signals that followed a more complex evolutionary history. One example is provided by the *TKL1* locus, which was shown to trigger considerable developmental differences in the brain when the modern or archaic human alleles are deployed ^34^, but that reached fixation of the derived allele in humans well before the Split (**Table S1**, chrX:153500000-153550000 HumanTMRCA ∼1200 kya).

As a perspective, given the current limitation in accurately dating derived substitutions fixed across both modern and archaic lineages, the only major events that could be pinpointed as defining the human lineage are the chromosome 2 fusion and the PAR2 translocation. Both events date back over 650 kya, predating the modern human, Neanderthal, and Denisovan divergence, and in this sense, they define all of us. Following these events, relaxation of selection due to reduced effective population size in the archaic populations, as well as the accumulation of cultural innovations in both modern and archaics, likely became the main drivers underlying the biological and behavioural differences observed in the archaeological record between what we can consider different populations of a single, cosmopolitan species.

## Material and Methods

### Genomic Data

Publicly available Whole genome sequences (WGS) for modern humans were obtained from https://ftp.1000genomes.ebi.ac.uk/vol1/ftp/release/20130502/, WGS for the archaic Neanderthal and Denisova and for the SS6004480 human Chromosome X were obtained from http://cdna.eva.mpg.de/neandertal/altai/Denisovan/ ; http://cdna.eva.mpg.de/neandertal/altai/AltaiNeandertal/VCF/ ; http://cdna.eva.mpg.de/neandertal/Spy94a/; http://cdna.eva.mpg.de/denisova/Den8/; http://cdna.eva.mpg.de/neandertal/altai/ModernHumans/bam/ ; http://cdna.eva.mpg.de/neandertal/Vindija/VCF/Vindija33.19/. Genotype calls for the PAR2 were obtained from https://s3-us-west-2.amazonaws.com/human-pangenomics/index.html?prefix=T2T/CHM13/assemblies/variants/1000_Genomes_Project/chm13v2.0/ PAR2 sites were mapped on the T2T-CHM13v2.0 reference. Genomic markers were instead mapped on build hg19 and analyses were run only at sites where the ancestral allele inferred from the 1000 Genomes project was available.

### Computing window-based genomic divergences

We computed genetic divergences from the human reference sequence (hg19) by counting the number of differences between the latter and Altai Neanderthal, Denisova or 50 Sub-Saharan African individuals from the 1000 Genomes Project over a sliding window of 50 kb across all the autosomes and chromosome X. We used phased haploid sequences for modern humans and picked a random allele at heterozygous sites in archaic humans. We then converted the resulting divergence estimates ((number of differences/2) / sequence length) in coalescence dates (or time since the most recent common ancestor (TMRCA)) by dividing it by the average human mutation rate (1.25 x 10^-8^ mut/bp/gen )^35^ and multiplying it by 30 years, here taken as the human generation time.

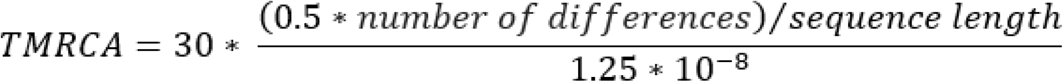

For each genomic window, we thus obtained a point estimate of the divergence between each archaic human and the reference genome (used as a proxy for a random modern human). We then took the top 90% most divergent estimate based on the 50 African individuals of our dataset as a measure of the within-modern human divergence (humanTMRCA). It is worth noting that we based our humanTMRCA estimates on African genomes to minimize the impact of the Neanderthal introgression that occurred after the expansion of modern humans out of Africa about 60 kya^20,36^. Neanderthal introgressed regions embedded within the reference genome are expected to yield deeper divergences between all tested Africans and the reference genome, but we assumed they would have a marginal impact overall, irrelevant for our experimental design aimed at detecting regions with increased similarity between modern humans and Neanderthals. Eventually, we filtered out from our downstream analyses regions with <50% of accessible sequence based on the 1000 Genomes Project strict accessibility masks (**Table S1**) and <90% availability of archaic and ancestral alleles along all 50 kb sites.

### Coalescent simulations to validate the obtained TMRCAs

To assess the accuracy of TMRCA estimates based on 50-kb windows, we simulated sequences that had experienced complete selective sweeps 500,000 years ago and inferred their TMRCA based on 50-kb windows.

We used msprime^37^ to generate 10 diploid chromosomes of length 5 Mbp for 51 individuals, assuming a constant recombination rate of 1.25×10^-8^ rec/bp/gen, a mutation rate of 1.25×10^-8^ mut/bp/gen, a generation time of 30 years, and a constant population size of 15,000 individuals. For each 50-kb window, we simulated a complete selective sweep 500,000 years ago, targeting the middle SNP (using SweepGenicSelection, with a start frequency of 1/2Ne, an end frequency > (2Ne - 1) / (2Ne) (here, 0.999999), a selection coefficient (s) of 1, and a time increment before the sweep (dt) of 1×10^-6^). We used one individual as a reference genome to define the genotypes of the 50 remaining individuals as reference or alternative alleles.

We then estimated the TMRCA for each 50-kb window along the genome, following the approach described above.

### Definition of Human650 and WMTH regions

Based on the data available in **Table S1**, we defined Human650 regions as regions with a) a HumanTMRCA <= 650 kya (i.e., after the separation of our lineage from that of archaic humans, referred to throughout the paper as the “Split”) and b) a DenisovaTMRCA >=650 kya (the date range expected from the population tree, Fig. 1A), to exclude genomic regions that experienced apparent relaxation of the molecular clock due to purifying selection.

Of these Human650 regions, we further identified What Made Them Human (WMTH) regions as those with i) a NeanderthalTMRCA <=650 kya (not expected on the basis of the population tree, unless human-Neanderthal admixture is involved), ii) at least one fixed derived mutation shared between Altai Neanderthal and modern humans (derived frequency >=0.98 in 1000 Genomes Project individuals) but not Denisovans and iii) no fixed derived mutations shared between Denisovans and modern humans but not Altai Neanderthal.

### Enrichment analysis

We explored the distribution of archaic/modern and modern/modern TMRCAs computed in **Table S1**, looking for enrichments of Neanderthal/human TMRCA compared to Denisova/human TMRCAs within autosomal loci, hence excluding the X chromosome from this particular analysis. We first examined regions displaying an archaic/human TMRCA falling within a moving cut-off (t) from 1000 to 100 kya, with a step of 10 ky and counted the number of 50 kb regions with a NeanderthalTMRCA less than *t* years ago and with DenisovaTMRCA>=650 kya, labelled as NRDA (Neanderthal Recent, Denisova Ancient), then we estimated the number of segments with DenisovaTMRCA less than t years ago and with NeanderthalTMRCA >=650 kya, these segments were labelled as DRNA (Denisova Recent - Neanderthal Ancient). We then tested whether the abundance of NRDA and DRNA regions was merely due to Incomplete Lineage Sorting by computing the normalized ratio:

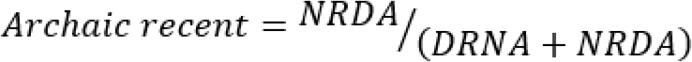

with

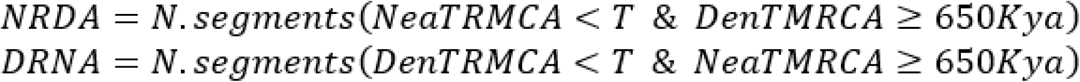

(Blue line in **Supplementary Figure S2)**. A value higher than 0.5 means an excess of NRDA regions compared to DRNA regions.

We then computed the same ratio by looking only within Human650 regions and obtained the Archaic Recent within Human650 enrichment curve (Green in **Supplementary Figure 2**).

### Relate

For each 50kb autosomal WMTH region, we further delved into the analysis of their TMRCA using Relate v1.1.9 ^38^. Trees were built for each region of interest using the entire 1000 Genomes dataset (3202 samples) and three archaic genomes. With this, we inferred genomic regions corresponding to each tree (regions with no inferred historical recombination within the sample).

We then computed the TMRCA of each tree spanning at least 5kb using the procedure described above (**Supplementary Figure S3**).

### Variant Annotation and analyses

We used Ensembl VEP (Ensembl GRCh37 release 113)^39^ to identify the most severe consequence and the gene associated with each site fixed derived in at least 98% of all 1000 Genomes Project individuals and falling in genomic windows with a HumanTMRCA <=1000 kya (i.e., that experienced a human fixation event after Event 1). We subsequently categorized consequences into three main classes:

- Coding (missense_variant, splice_acceptor_variant, splice_donor_5th_base_variant, splice_donor_region_variant, splice_donor_variant, splice_polypyrimidine_tract_variant, splice_region_variant, start_lost, stop_gained, stop_lost)
- Regulatory (3_prime_UTR_variant, 5_prime_UTR_variant, mature_miRNA_variant, regulatory_region_variant, TF_binding_site_variant)
- Others (downstream_gene_variant, intergenic_variant, intron_variant, non_coding_transcript_exon_variant, synonymous_variant, upstream_gene_variant)

We referred to Coding and Regulatory consequences together as Functional and distinguished them into four categories: FuncH (>=98% derived allele frequency only in modern humans), FuncHD (>=98% derived allele frequency in modern humans and homozygous derived in Denisova but ancestral in Altai Neanderthal), FuncHN (>=98% derived allele frequency in modern humans and homozygous derived Neanderthals but ancestral in Denisova), and FuncHH (>=98% derived allele frequency in modern humans and homozygous derived in Altai Neanderthal and Denisova).

We then divided the Functional sites into non-overlapping sets based on the HumanTMRCA and ArchaicTMRCA displayed by the 50 kb windows containing them:

Human350: HumanTMRCAs <=350 kya, DenisovaTMRCA >650 kya;
Human650 350 kya < HumanTMRCA <= 650 kya and DenisovaTMRCA > 650 kya;
A final group, called WMTH, was formed by a subset of Human350 regions displaying a NeanderthalTMRCA <= 650 kya.

To investigate the biological network formed by genes having undergone functional modifications in the past 650 ky of modern human evolution, those affected by at least one FuncH or FuncHN variant of the Human350 and Human650 sets were analyzed using the STRING v12.0 database (STRING database https://string-db.org/). The resulting network of protein-protein interactions was analysed for phenotype enrichment using the STRING enrichment tool against the Monarch Initiative database^40^.

### PAR 2 analyses

We obtained information about sequencing depth along the X chromosomes of a Neanderthal (Spy), a Denisova (Denisova 8), and a modern human male (SS6004480) individual by computing the average depth over non-overlapping 10 kb long windows along the whole chromosome X and displayed the distribution of sequencing depth for the PAR1 (X:0-2,394,410), PAR2 (chrX:153,925,834-154,259,566)^41^, and remaining portion (chrX:2,394,411-153,925,833) of the X chromosome. To graphically compare the coverage level between the reference modern human, Neanderthal, and Denisova for each of these regions, we scaled it using the *scale* function in R and adding 1 to each value separately.

As PAR2 behaves similarly to an autosomal region, it is not possible to distinguish short reads coming from the X-linked from Y-linked PAR2 sequence in a given genome. We therefore retrieved all PAR2-derived reads in modern human male individuals mapping onto the X chromosome, using the most up-to-date set of genotype calls made available by Rhie et al.^41^ for the 1000 Genomes Project panel^42^. In practice, we exploited the genotype calls performed by the T2T Y chromosome consortium on all 1000 Genomes Project samples by mapping their reads onto the latest T2T-CHM13v2.0 reference sequence after masking out the chromosome Y-PAR2 region. In this way, we let the X-PAR2 region (chrX:153925834-154259566, almost identical to the Y-PAR2 region chrY:62122809-62460029) attract the relevant reads from either chromosome in both males and females^41^.

In addition, we estimated allele frequencies for all non-reference PAR2 sites in 1000 Genomes Project modern human males and females separately. We used the observed allele frequencies in female and male X chromosomes to infer the allele frequency in male Y chromosomes with the following:

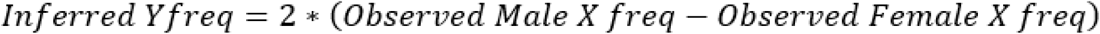

We identified 11 sites that reached an allele frequency in male Y chromosomes close to 1, hence distinguishing Y-PAR2 from X-PAR2 sequences. Other classes of notable mutations, namely the ones showing an X frequency of less than 0.5 in males (inferred to be less than 1 in Y-PAR2) and of approximately 0 in females, as well as the ones showing an X frequency of approximately 0.5 in males (inferred to be approximately 1 in PAR2-Y chromosomes) and of more than 0 in females, were annotated. However, we subsequently discarded them as respectively representing not yet fixed male-specific mutations or mutations present in the X-PAR2 at the time of the translocation, placing them outside of the scope of the present study. The 11 mutations found to be fixed in all surveyed males and virtually absent from females spanned 21,215 bps at the 5’ end of the Y-PAR2 sequence and were considered to have occurred since the divergence between X-PAR2 sequences and the most recent common ancestor of all Y-PAR2 sequences - in other words, before the diversification of present-day Y chromosome lineages. We then calculated the time taken for these mutations to accumulate by using the simple formula:

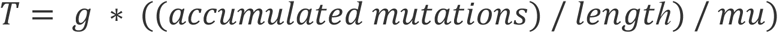

using 30 years as the generation time (g) in humans and ruling out complex back-and-forth scenarios between the X and Y PAR2 pools, given the absence of these mutations within the X-PAR2 gene pool. Using the Y chromosome mutation rate of 3 x 10^-8^ mut/bp/gen^32^ yielded a T of 518 ky (95% CI: 295-915 ky) before the MRCA of modern humans. We computed confidence intervals around our estimate by evaluating the probability distribution of observing exactly 11 mutations in 21,215 bp over different time intervals and taking the 2.5th and 97.5th quantiles of this distribution. Since the MRCA of modern humans has been attested to date back to338 kya, we eventually obtained a final point estimate of 856 kya as the time of the PAR2 translocation.

## Acknowledgements

The authors would like to thank Prof. Gil Guastoni Rosenthal, Dr. Andra Meneganzin, and Giacomo Moro Mauretto for the fruitful discussions during the drafting of the current manuscript, and Dr. Mathilde André for facilitating access to Papuan allele frequencies. LP was funded by the Italian Ministry of University and Research PRIN N. 2022B27XYM and N. P2022MZCNX.

## Data and materials availability

All genomic data are publicly available at https://ftp.1000genomes.ebi.ac.uk/vol1/ftp/release/20130502/, http://cdna.eva.mpg.de/neandertal/altai/Denisovan/; http://cdna.eva.mpg.de/neandertal/altai/AltaiNeandertal/VCF/; http://cdna.eva.mpg.de/neandertal/Spy94a/; http://cdna.eva.mpg.de/denisova/Den8/; http://cdna.eva.mpg.de/neandertal/Vindija/VCF/Vindija33.19/, TMRCA computations are available as supplementary data.

