## Supplementary Information for "Partitioning the genomic journey to becoming *Homo sapiens*"

*Affiliations*
*1 Department of Biology, University of Padova, Via Ugo Bassi 58/B, 35131, Padova, Italy*
*2 Institute of Genomics, University of Tartu, Riia 23/b, 51010 Tartu, Estonia*
*3 Bioinformatics Research Centre, Aarhus University, Universitetsbyen 81, 3., Building 1872, DK-8000 Aarhus C Denmark*
*4 Institute of Organismic and Molecular Evolution, Palaeogenetics Group, Johannes Gutenberg University, Mainz 55128, Germany*
*5 Department of Neurosciences, University of Turin, 10126, Turin, Italy*
*6 Istituto di Calcolo e Reti ad alta prestazione, CNR, 80131, Naples, Italy*
*7 Monash University Malaysia Genomics Facility, School of Science, Monash University, 47500 Bandar Sunway, Selangor Darul Ehsan, Malaysia*
*8 The Jackson Laboratory for Genomic Medicine, Farmington, CT, USA*

**Supplementary Information**

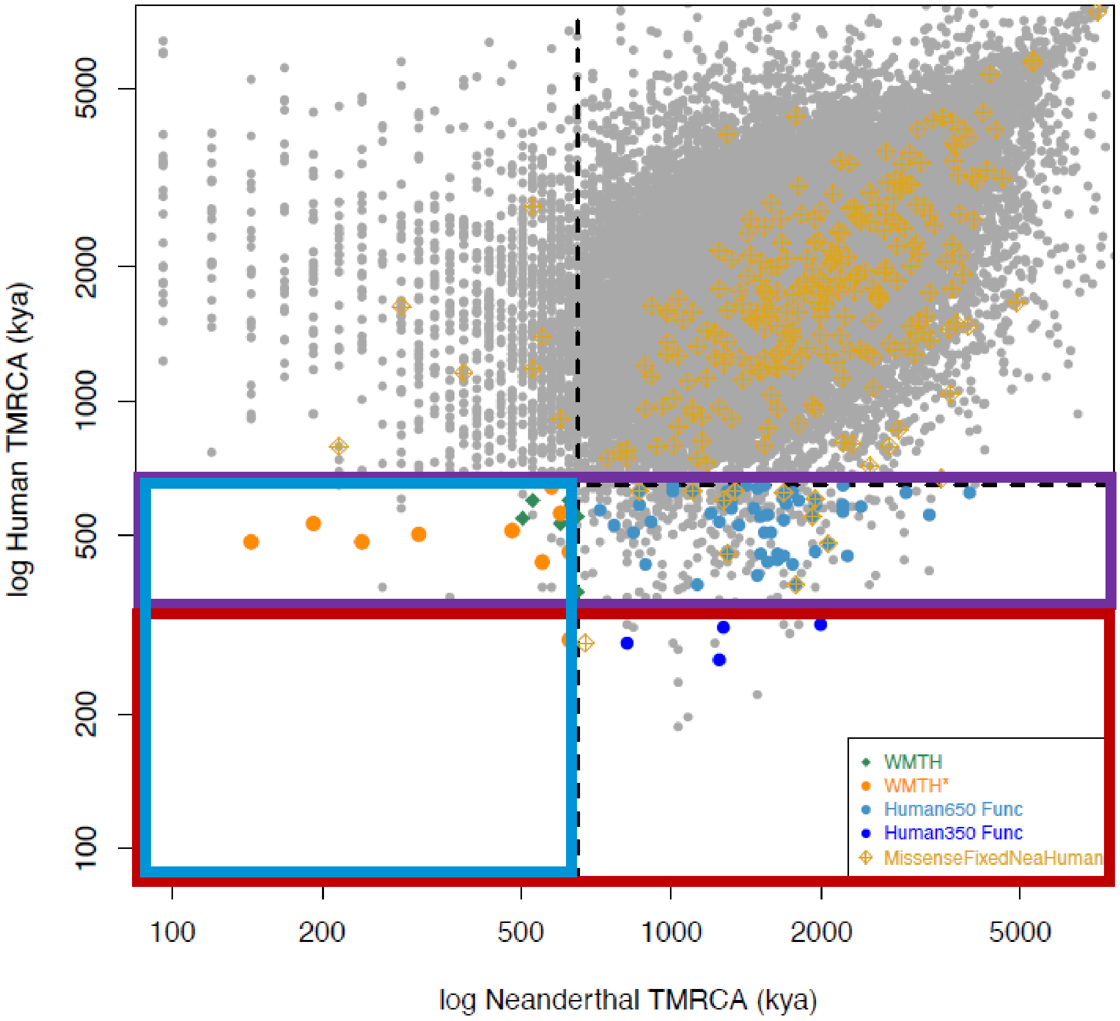


**Figure S1. Scatterplot of the Neanderthal-Human reference and the Human-Human TMRCAs on windows displaying a Denisova-Human reference TMRCA >=650 kya.** Grey dots represent the bulk of the distribution, which as expected is centered around the diagonal for x and y values >=650 kya. Dashed lines report the Split event (i.e., the separation of modern and archaic humans) along both axes. The top right quadrant appears enriched compared to the bottom left quadrant as an effect of the reported Neanderthal-modern human introgressions. The top left quadrant is, as expected, enriched by Neanderthal regions introgressed inside the reference sequence and this signal is assumed not to impact our results, since the resulting human-human TMRCA (y-axis) is above the thresholds used for our downstream analyses. Orange dots and green diamonds respectively correspond to the 50 kb WMTH regions reported in Table 1 with more (orange) and less (green) stringent criteria. Light blue dots represent 50 kb Human650 regions - that is, including at least one functional (coding or regulatory) variant with a human-derived frequency >=98% and absent in Denisovans and with a Human TMRCA <=650 kya (i.e., more recent than the Split). Dark blue dots, on the other hand, identify the subset of these regions with a Human TMRCA <=350 kya (more recent than the Introgression, Human350). Finally, golden diamonds flag regions containing at least one missense variant with a derived frequency of at least 98% in humans and fixed ancestral in both the Altai Neanderthal and Denisova, as reported in[1].

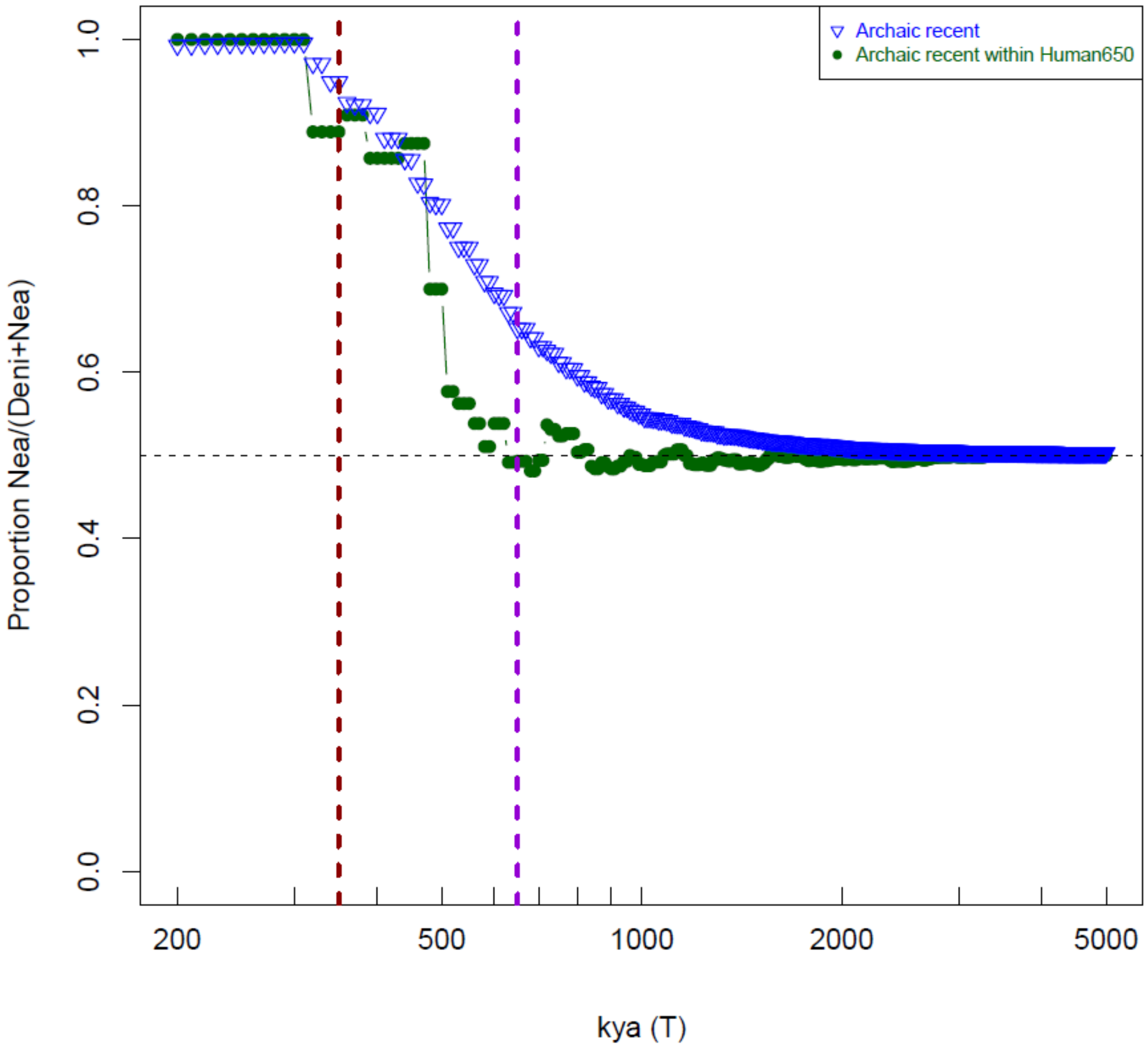


**Figure S2. Relative Archaic Proportion of the candidate regions.** The dark blue line (Archaic recent) reports the amount of Neanderthal regions coalescing with the hg19 human reference at time T while Denisova coalesceces with it >=650 kya (NRDA), divided by the sum of the latter and the number of regions with a Denisova-human reference coalescence at T and a Neanderthal-human reference coalescence dating back to >=650 kya (DRNA). The formula applied is NRDA/(DRNA+NRDA). A ratio of 0.5 means that the amount of NRDA regions is the same as that of DRNA regions.

The dark-green line (Archaic recent within Human650) represents the previous proportion, computed only for regions showing a coalescence event younger than 650 ky in all modern humans (Human650 regions). The vertical dotted lines indicate the Introgression (red) and the Split (violet).

**Relate analyses of WMTH regions**

For each 50 kb autosomal WMTH region (*Materials andMethods*), we further delved into the analysis of their TMRCAs using Relate v1.1.9[2].

We In practice, we corroborated our findings based on coalescence times estimated over fixed 50 kb windows by inferring *Relate* sub-trees on every autosomal region reported in Table 1, except for the two pairs of adjacent 50 kb windows on chromosomes 10 and 16, which were collapsed into two 100 kb windows. Using the complete 1000 Genomes dataset (3202 samples) as well as the archaic genomes of Denisova and Neanderthal, we then computed the TMRCA between modern humans and modern humans, Neanderthal, and Denisova for each *Relate* tree spanning >= 5kb (Figure S3).
All the regions for which our initial estimates provided a Human-Neanderthal TMRCA at <=350 kya displayed a *Relate* sub-tree consistent with the Denisova, (Neanderthal, Human) topology expected given our conditions (Figure S3, chromosomes 6,7 and 16). Regions with deeper Human-Neanderthal TMRCAs (Figure S3, chromosomes 1, 10 and 12), on the other hand, presented[2] a combination of trees with and without the expected features (outlined by the dashed lines in **Figure S3)**, consistent with recombination events having occurred after the admixture event within each 50 kb windows considered in our analyses.

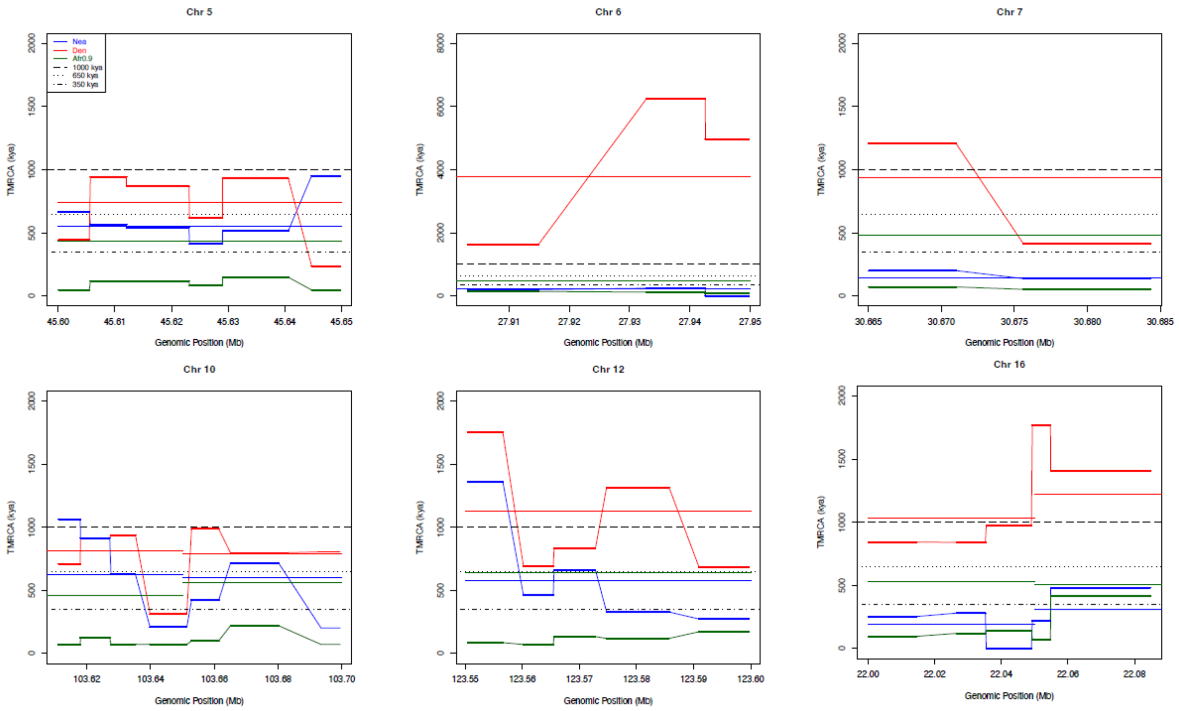


**Figure S3. TMRCA of *Relate* trees for each candidate genomic region.** Continuous thin lines represent the average TMRCA estimated for each whole 50 kb region under consideration (green for Africans, blue for Neanderthals, red for Denisova) and reported in Table 1. The thick line represents the estimates for each *Relate* tree longer than 5kb identified within said region. Dotted lines represent the time thresholds of 650 kya (Split) and 350 kya (Introgression). The TMRCAs associated with the sub-trees of the regions falling on chromosomes 6, 7 and 16 fit within the expected scenario, while recombination events yield a more composite scenario for the other regions. Abbreviations: Afr0.9, top 90% deepest TMRCA between 50 African individuals and the human reference; Nea, Altai Neanderthal - human reference TMRCA; Den, Denisova - human reference TMRCA.

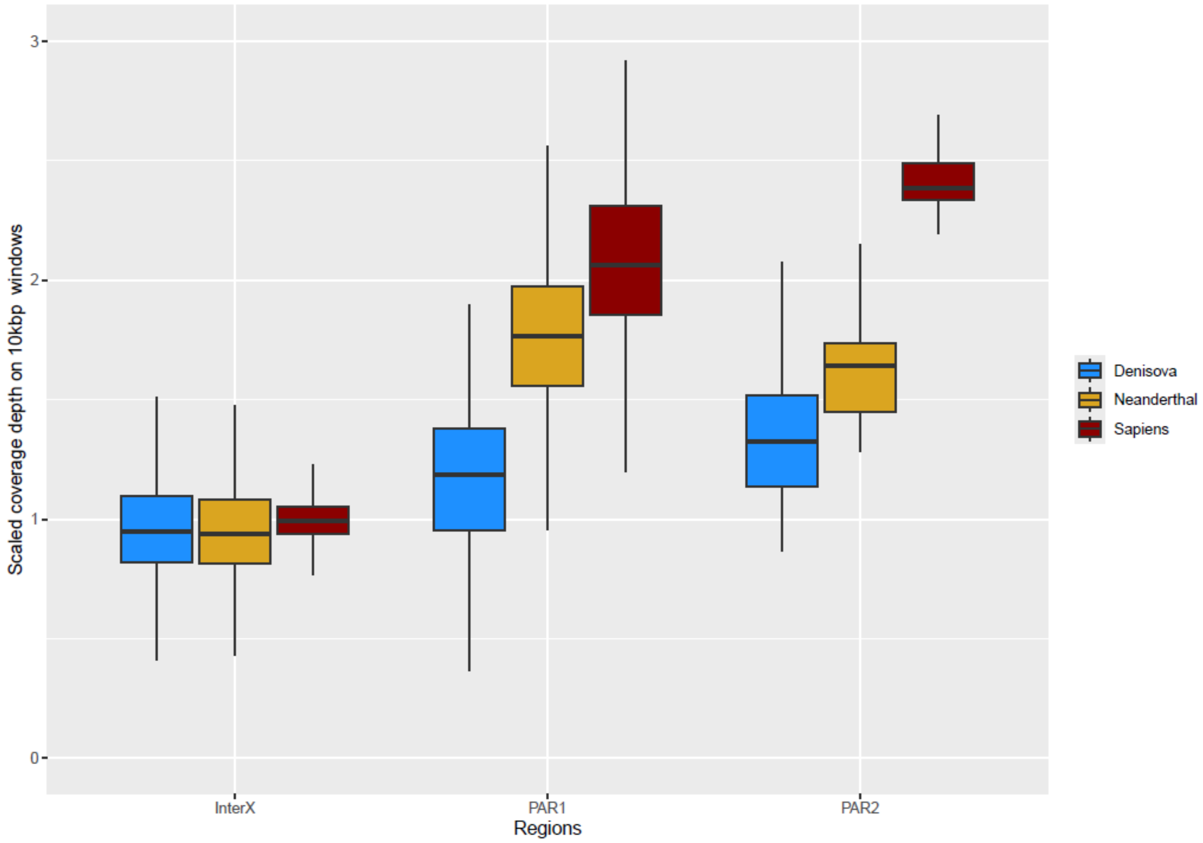


**Figure S4.PAR2 coverage depth**

Spy Neanderthal, Denisova and Modern human (SS6004480 individual) sequencing depth averaged over windows of 10 kb from the central part of chromosome X, X-PAR1 and X-PAR2. The sequencing depths displayed are z-transformed based on the depth distribution of the internal part of the relevant chromosome X (InterX). In all modern and archaic genomes, both PAR regions show a statistically significant (Mann-Whitney p-value<1.6 x 10-6) higher depth compared to their InterX, consistent with the presence of both a Y and an X chromosome PAR sequences in the sampled individuals. Y reads are expected to map on the X chromosome because of the high sequence homology at PAR2 sites.
